# 3D genome organization drives gene expression in trypanosomes

**DOI:** 10.1101/2023.04.01.535209

**Authors:** Florencia Díaz-Viraqué, María Laura Chiribao, Gabriela Libisch, Carlos Robello

## Abstract

In trypanosomes —eukaryotic unicellular pathogens that cause disabling human and animal diseases— very few transcriptional regulatory elements have been described and it is largely accepted that they regulate gene expression mainly post-transcriptionally. In this regard, the role of the spatial organization of the genome on gene expression and vice versa remains practically unexplored. The genome of these parasites is partitioned into core (highly conserved syntenic) and species-specific disruptive regions (synteny disruption), containing multigene families encoding for surface glycoproteins. By mapping genome-wide chromatin interactions we demonstrate that these regions constitute 3D compartments (C and D). These chromatin compartments present significant differences in DNA methylation, nucleosome positioning and chromatin interactions, affecting genome expression dynamics. We show that the genome is organized into chromatin folding domains and transcription is dramatically determined by the local chromatin structure. Our results support a model in which epigenetic mechanisms dramatically impact gene expression in these eukaryotic pathogens.

## Main

Gene organization, regulation of genome expression and RNA metabolism in trypanosomes present some characteristics that distinguish them from other eukaryotes. In these organisms, genes are organized into directional gene clusters (DGCs) separated by short sequences denominated strand-switch regions (SSRs), where the transcription sense converges or diverges. Although sequence-specific promoters for RNA polymerase II were recently reported (Cordon-Obras et al., 2022), classical enhancers have not been identified. The DGCs are transcribed as large polycistronic transcription units (PTUs) (Johnson et al., 1987), and messenger RNA maturation implies co-transcriptional trans-splicing of spliced leader RNA and polyadenylation (Matthews et al., 1994). Consequently, post-transcriptional regulation has been proposed as the main gene expression regulation level (Clayton et al. 2007 and 2019; Chávez et al., 2021). However, it is not sufficient to explain the critical regulatory changes necessary for pathogenicity, and recent reports have highlighted the relevance of histone posttranslational modifications, histone variants, base J, nucleosome positioning, and chromatin organization on the regulation of gene expression, cell cycle control, differentiation, and pathogenesis (Maree et al., 2022; Lima et al., 2021 and 2022; Faria et al., 2021; Respuela et al., 2008; Rosón et al., 2022; Nunes et al., 2020; Ramos et al.,2015).

Epigenetic mechanisms have been described as a key factor in transcriptional control in other protozoan parasites (Lizarraga et al., 2020; Bunnik et al., 2018). In trypanosomes, 5-methylcytosine (5mC) modification mark was demonstrated in nuclear DNA (Rojas et al., 1990; Militello et al., 2008), while the presence of N6-methyladenine (6mA) was suggested but not evidenced in *T. cruzi* (Rojas et al., 1990). In addition, a functional role of 5mC methylation was suggested in *T. cruzi* through incubation with 5-azacytidine (Rojas et al., 1991). However, the genomic distribution of these DNA methylation marks and its possible role in controlling gene expression and pathogenesis remain unexplored. Recently, it has been shown that a specific genome organization is required for monogenic expression of the Variant Surface Glycoprotein (VSG) expressed on *T. brucei* surface (Faria et al., 2021), denoting the importance of the genome architecture on gene expression in trypanosomes. Despite this, the role of chromatin organization controlling gene expression of non-antigenic genes and gene expression and pathogenesis in *T. cruzi* remains practically unexplored.

The surface coat of trypanosomes is covered by many proteins (Ferguson 1997). Nevertheless, its composition and expression vary among different species, reflecting differences in their life cycles and probably the molecular mechanisms that underlie the regulation of the expression of these genes. *T. brucei* remains extracellular during its life cycle, and the efficient humoral immune evasion relies on antigenic variation, which systematically alters the VSG protein displayed to the host immune system (Vickerman K 1969; Cross GAM 1975). On the other hand, *T. cruzi*, whose life cycle has an intracellular and an extracellular stage, uses antigenic variability, simultaneously expressing thousands of slightly different surface proteins (Buscaglia et al., 2004; dos Santos et al., 2012; De Pablos et al., 2012). Genes encoding surface proteins are found in specific locations of the genome. The genomes of trypanosomes have been described as being partitioned into two large regions previously named compartments (Berná et al., 2018; Müller et al., 2018). In *T. cruzi*, the core compartment (which presents synteny among trypanosomatids) is composed of conserved genes, while the disruptive (loss of synteny) compartment is where most of the surface coding genes are located; in particular, the disruptive compartment is enriched in mucins (TcMUC), mucin-associated surface protein (MASP), and trans-sialidases (TS) coding genes (Berná et al., 2018). In *T. brucei*, genes encoding for VSG surface proteins are also located in a particular compartment in the genome, which is located mostly in the extreme of the chromosomes, and it is generically named subtelomeric (Müller et al., 2018).

Stage-specific expression of surface proteins is essential for the life cycle of trypanosomes; therefore, this large group of genes needs to be tightly regulated and physical separation in the genome could be an efficient mechanism to regulate its expression. Based on these observations, the central hypothesis of this paper is that this specialized genome organization enables different regulation mechanisms to occur, where chromatin structure plays a central role as the first level of gene expression regulation in trypanosomes. To test this hypothesis and better understand the gene expression regulatory mechanisms underlying the infection process, we explored how the spatial organization of the genome influences gene expression in the context of previously defined genome compartments. Our results provide novel insights into the connection between genome organization and gene expression.

## Results

### The genome of trypanosomes is organized into chromatin-folding domains

Based on our previous observation that the *T. cruzi* genome is partitioned into core and disruptive compartments (Berná et al., 2018), we wondered whether this organization correlates with its three-dimensional organization. For this analysis, we classified the chromosomes into core (>80% core compartment), disruptive (>80% disruptive compartment), and mixed (Supplementary Table 1). To interrogate the chromatin conformation in *T. cruzi* cells, we mapped genome-wide interactions using Chromosome conformation capture (Hi-C) data from epimastigotes previously used for genome assembly (Wang et al., 2021). We evaluated the interaction matrices at a variety of bin sizes, and the analysis of the data revealed that there are several regions in close spatial proximity indicating the presence of chromatin folding conformations (Fig. 1a). To identify all such topological domains in the genome, we explored the formation of chromatin folding domains by using three different algorithms (Supplementary Table 2 and Supplementary Fig. 1). As in other organisms (Fraser et al., 2015), the structural domains are organized in a hierarchical way showing that there are different levels of the organization. Using a 5 kb resolution Hi-C map, 173 domains with 129 kb mean size were identified, occupying 60% of the genome. We noticed that a considerable fraction of the genome remains structured in these folding domains representing a prominent feature of genome organization. However, the structures identified are smaller than Topologically Associating Domains (TADs) described in other eukaryotes (880 kb mean size of topological domains in the mouse stem cell genome) (Dixon et al., 2012). In this sense, TAD predictor algorithms fail to identify domains in the smallest chromosomes (< 1.1 Mbp) and at common resolutions for the identification of TADs. This makes sense since the chromosomes of *T. cruzi* are smaller (less than 3 Mbp). For this reason, we prefer to refer to them as chromatin folding domains (CFDs).

**Fig. 1:**
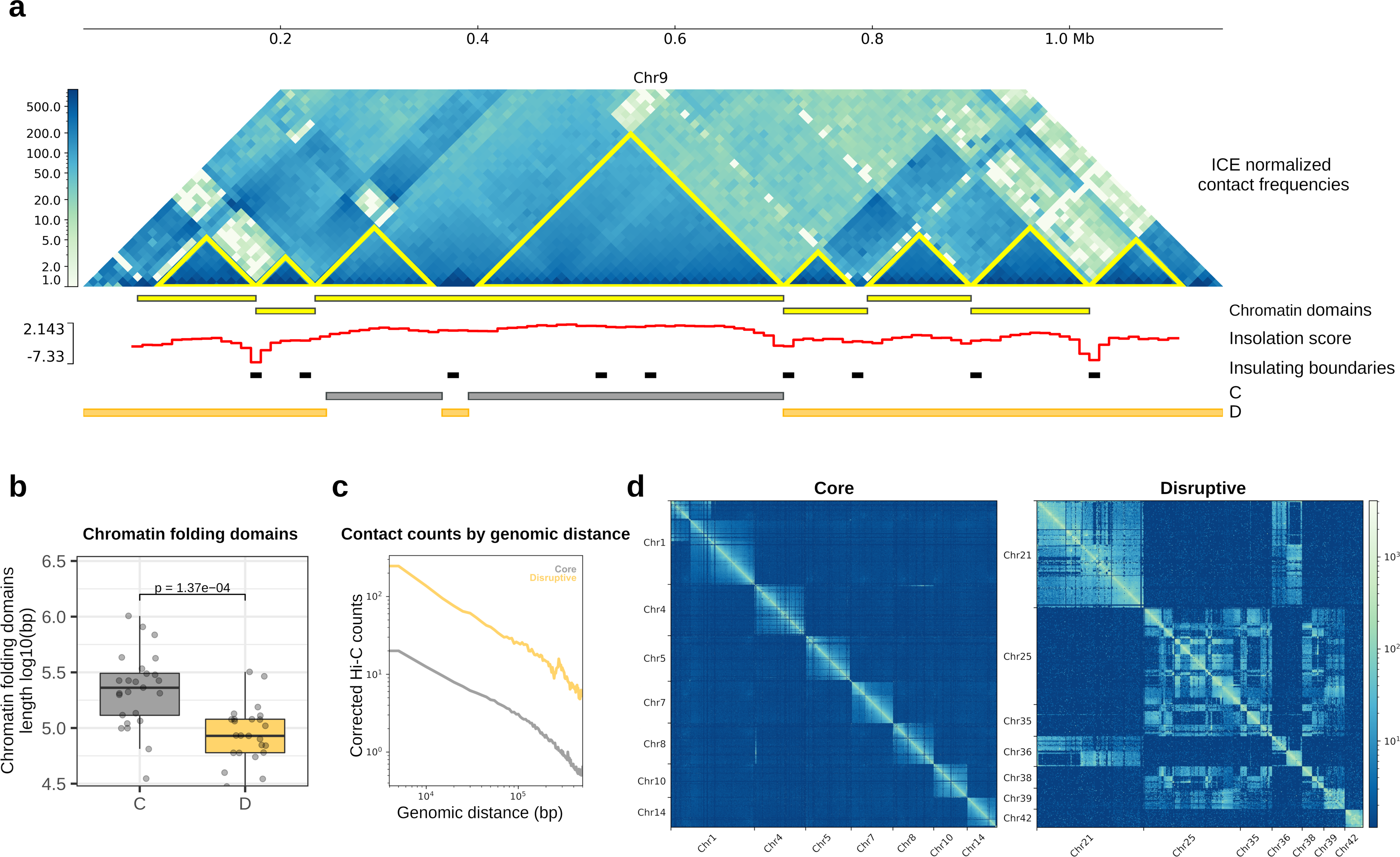
Chromatin interaction in *T. cruzi* genome compartments. **a,** Normalized Hi-C interaction frequencies of a representative chromosome (chromosome 9) displayed as a two-dimensional heatmap at 10 kb resolution. Chromatin Folding Domains (CFDs) identified using HiCExplorer and TADtools (triangles on Hi-C map) are indicated in yellow. There is a correlation between the local minimum of insulation score calculated using FAN-C, which represents the region between two self-interacting domains, and the CFDs identified with HiCExplorer and TADtools. The genomic position of core and disruptive genome compartments are represented in grey and yellow, respectively. **b,** Boxplot of CFD length, depending if the domain is in the core or disruptive compartment. CFD core median length = 230 kb (N = 25), CFD disruptive median length = 85 kb (N = 24). Significance was determined using an Unpaired two-samples Wilcoxon test (two-sided), *p-value* = 1.37 x 10^-4^. **c,** Mean interaction frequencies at all genomic distances at 5 kb resolution. **d,** Intra- and inter-chromosomal interactions between core and disruptive chromosomes. C: core, D: Disruptive.

We analyzed the self-interacting chromosomal domains in the context of the core and disruptive genome compartments, and we observed that both present CFDs. Notably, 80% of the identified domains belong to only one compartment, indicating that CFDs in each compartment are mutually exclusive (Supplementary Table 2). Therefore, the previously defined linear genome compartments (Berná et al., 2018) correspond to three-dimensional compartments of the nucleus. In addition, we found that the disruptive compartment exhibits smaller CFDs (Fig. 1b) and a higher frequency of interactions at different distances (Fig. 1c). We next investigated the chromosomal interactions and found that the disruptive compartment exhibits more inter-chromosomal contacts. In contrast, the interactions in the core compartment are mainly intra-chromosomal (Fig. 1d). Finally, we analyzed the chromatin isolation regions to determine if specific genes could act as isolators; of the 140 genes detected, we found an enrichment in surface genes and transposable elements (Supplementary Table 3).

### Compartmentalized regions of the genome exhibit different patterns of gene expression

As chromatin folding domains are defined as genomic regions with abundant self-self contacts and fewer contacts with surrounding areas (Dixon et al., 2016), the 3D organization of the genome is expected to exhibit global implications in gene expression. To explore this, we performed RNA-seq of different stages of the parasite life cycle (epimastigotes, intracellular amastigotes and cell-derived trypomastigotes) and genes were classified into high, medium, and low expressed genes according to the number of transcripts detected. We found that the core compartment is mainly composed of moderate and high-expressed genes in all the stages analyzed. In contrast, the disruptive compartment, which in some cases comprises almost entire chromosomes, is enriched in low expressed genes or genes with undetectable RNA levels (Fig. 2a). This difference in expression between core and disruptive compartments is statistically significant in all the stages (Fig. 2b). Moreover, the core compartment exhibits similar levels of expression whereas mean expression of the disruptive compartment increases in trypomastigotes (Welch Two Sample t-test *p* value 4.59e-4 and 9.92e-4 in comparison with amastigotes and epimastigotes, respectively). To further investigate the expression profiles in the compartments, we examined the RNA levels of each gene. While core genes are expressed in similar proportions in all stages, the disruptive genes present lower overall expression (Fig. 2c, d and Supplementary Fig. 2), and bimodal distribution in the trypomastigote stage, where a discrete number of genes increase their expression (Fig 2d and Supplementary Fig. 2). Furthermore, differences in expression between core and disruptive compartments were also obtained with the Brazil A4 strain (Wang et al., 2021) (Supplementary Fig. 3). Hence, these differences in gene expression between genomic compartments are characteristic of *T. cruzi* and are not strain specific.

**Fig. 2:**
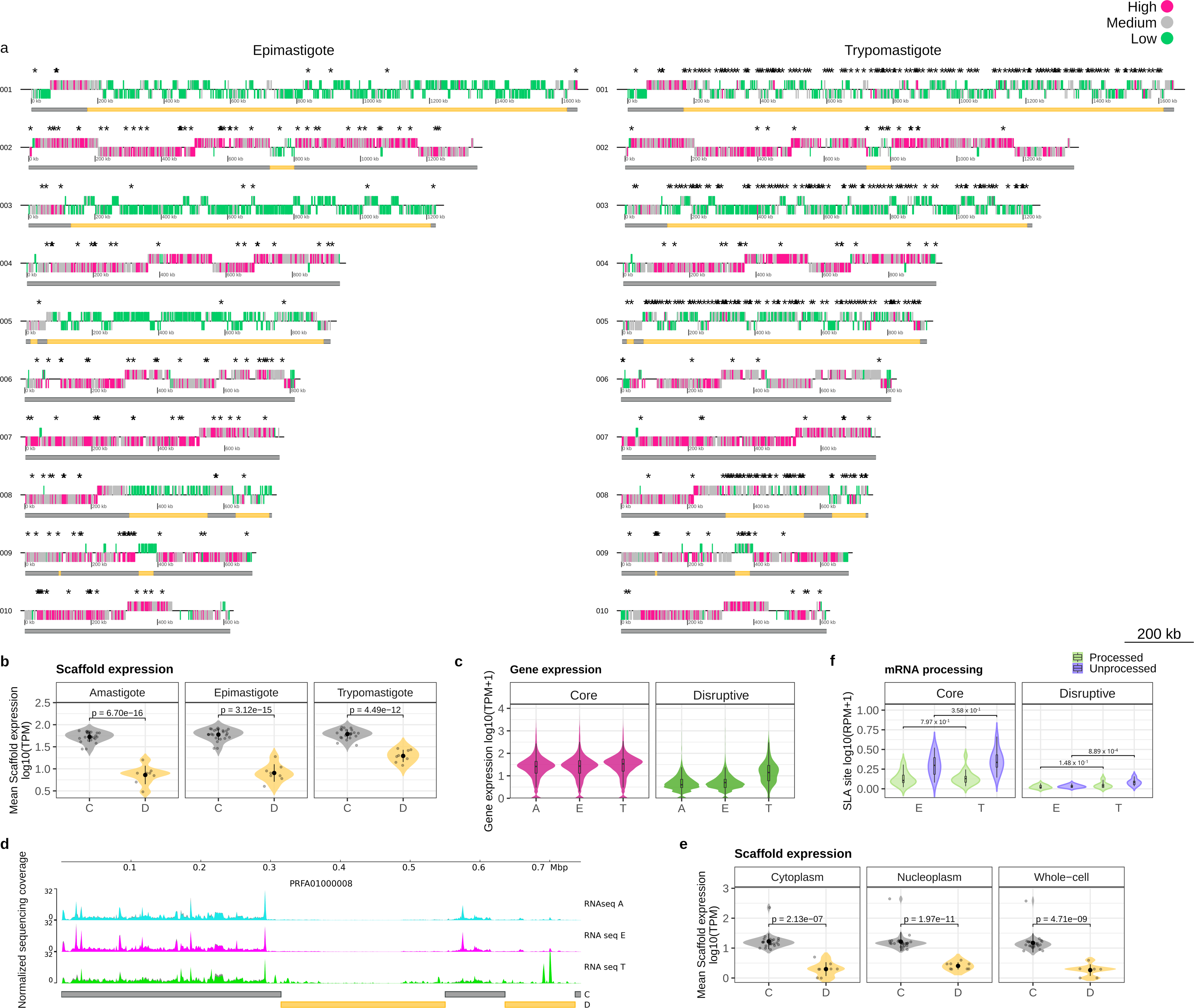
Transcriptional heterogeneity in the genomic compartments of *T. cruzi*. **a,** Genomic distribution of genes classified according to the number of transcripts in the largest ten scaffolds. Core and disruptive genomic compartments are indicated in gray and yellow, respectively. Differentially expressed genes are indicated as asterisks. **b,** Mean scaffold expression of scaffold composed by core or disruptive compartment. Scaffolds were classified as core or disruptive when one of the genome compartments spans 80-100% length of the scaffold. Scaffolds containing the core compartment present more mean expression than the scaffolds defined as disruptive. Values are *p-value* from Unpaired Welch Two Sample t-test. **c,** RNA expression of core and disruptive genes at the different stages of the parasite life cycle. The bimodality coefficient obtained for expression data of disruptive genes in trypomastigotes was 0.678. Values larger than 0.555 indicate the bimodality of data (Pfister et al., 2013). Mean expression (log10(TPM)) of core genes (N = 11947) are 1.41, 1.43 and 1.49 in amastigotes, epimastigotes and trypomastigotes, respectively. Mean expression (log10(TPM)) of disruptive genes (N = 2991) are 0.59, 0.64 and 1.17 in amastigotes, epimastigotes and trypomastigotes, respectively. **d,** Representative coverage plot normalized using counts per million (CPM). Bin size 10 bp. For each condition, two biological replicates are plotted. The genomic position of core (C) and disruptive (D) genome compartments are represented in grey and yellow, respectively. **e,** Mean scaffold expression of core and disruptive scaffolds using transcriptomic data of different subcellular compartments. Values are *p* value from Unpaired Welch Two Sample t-test. **f,** Quantification of the processed and unprocessed transcripts from RNA-seq reads covering the SLA site. SLA regions with more reads were selected for the analysis. Core processed *p* value = 0.797; Core Unprocessed *p* value = 0.358; Disruptive processed = 0.148; Disruptive unprocessed *p* value = 8.89 x 10^-4^. The *p* values are from the Wilcox test for the difference in means in all cases. C: core, D: Disruptive.

We next determined which genes were differentially expressed between the different stages. We found that 94% of genes enriched in epimastigotes belong to the core genomic compartment, while in trypomastigotes, 20% of differentially expressed genes (DEGs) are in the core, and 80% are in the disruptive compartment (Supplementary Tables 4 and 5). Thereby, the core compartment presents stage-specific genes in both stages, whereas the disruptive is a compartment characteristic of the trypomastigote stage, where long stretches of the genome concentrate many genes that are differentially expressed (Fig. 2a).

We investigated whether the reduced expression of the disruptive compartment in epimastigotes is due to transcriptional regulation or is mainly controlled by post-transcriptional mechanisms, which is widely accepted in these organisms. Given that post-transcriptional regulation mechanisms commonly occur in the cytoplasm and during nuclear exportation, we compared the expression patterns in cytoplasmic and nuclear transcriptomes (Pastro et al., 2017). We found that the differences of expression between compartments previously observed can already be observed in the nuclear transcriptome (Fig. 2e and Supplementary Fig. 4). Since these results provide evidence that the reduced expression of the disruptive compartment results from nuclear mechanisms of regulation of expression, we analyzed nascent mRNAs in the different stages taking advantage of the characteristic of the transcript maturation process in trypanosomatids. Due to the trans-splicing mechanism to produce mature mRNAs in trypanosomatids, it is possible to measure unprocessed transcripts from RNA-seq reads spanning the spliced leader acceptor (SLA) site, allowing us to quantify nascent or immature transcripts (López-Escobar et al., 2022). Our results showed the presence of similar levels of unprocessed transcripts corresponding to the core compartment in both stages and a reduction in the number of unprocessed transcripts from the disruptive compartment in epimastigotes, indicating reduced transcription or RNA maturation in this stage (Fig. 2f and supplementary Fig. S4). Taken together, our results confirm that the reduced expression of the disruptive compartment is a result of nuclear mechanisms of regulation.

### Well-positioned nucleosomes are enriched in the disruptive genome compartment

The nucleosome is the basic structural unit of DNA packaging in eukaryotes (Luger et al. 1997; Khorasanizadeh, 2004; Ammar et al., 2012), and nucleosome positions have been shown to be correlated with the transcriptional state (Workman, J. L. 2006). To test if genome organization determines the selective different expression patterns of compartments in a stage-specific way, we evaluated nucleosome positioning using MNase-seq data from epimastigotes and cell-derived trypomastigotes (Supplementary Table 6) (Lima et al 2021). We found that the core compartment exhibits a higher density of nucleosome occupancy in both stages, which is in concordance with previous reports (Lima et al., 2022), (Fig. 3a). However, we observed in our analyses that there is a correlation between the percentage of disruptive genes present in a scaffold and the density of well-positioned nucleosomes (Fig. 3b). We then analyzed the density of well-positioned nucleosomes and found that it was double in the disruptive compartment than in the core (Fig. 3c and d). To characterize the dynamic organization of nucleosomes, we compared nucleosome architecture local changes between epimastigotes and trypomastigotes in the compartments. We found that the changes in the nucleosome map were more prominent in the core compartment (Fig. 3e); there were few differences in the presence/absence of nucleosomes between stages, and nucleosome shifting is the most abundant alteration. On the contrary, few changes were detected in the disruptive compartment, results that are consistent with more static regions regarding nucleosome movement.

**Fig. 3:**
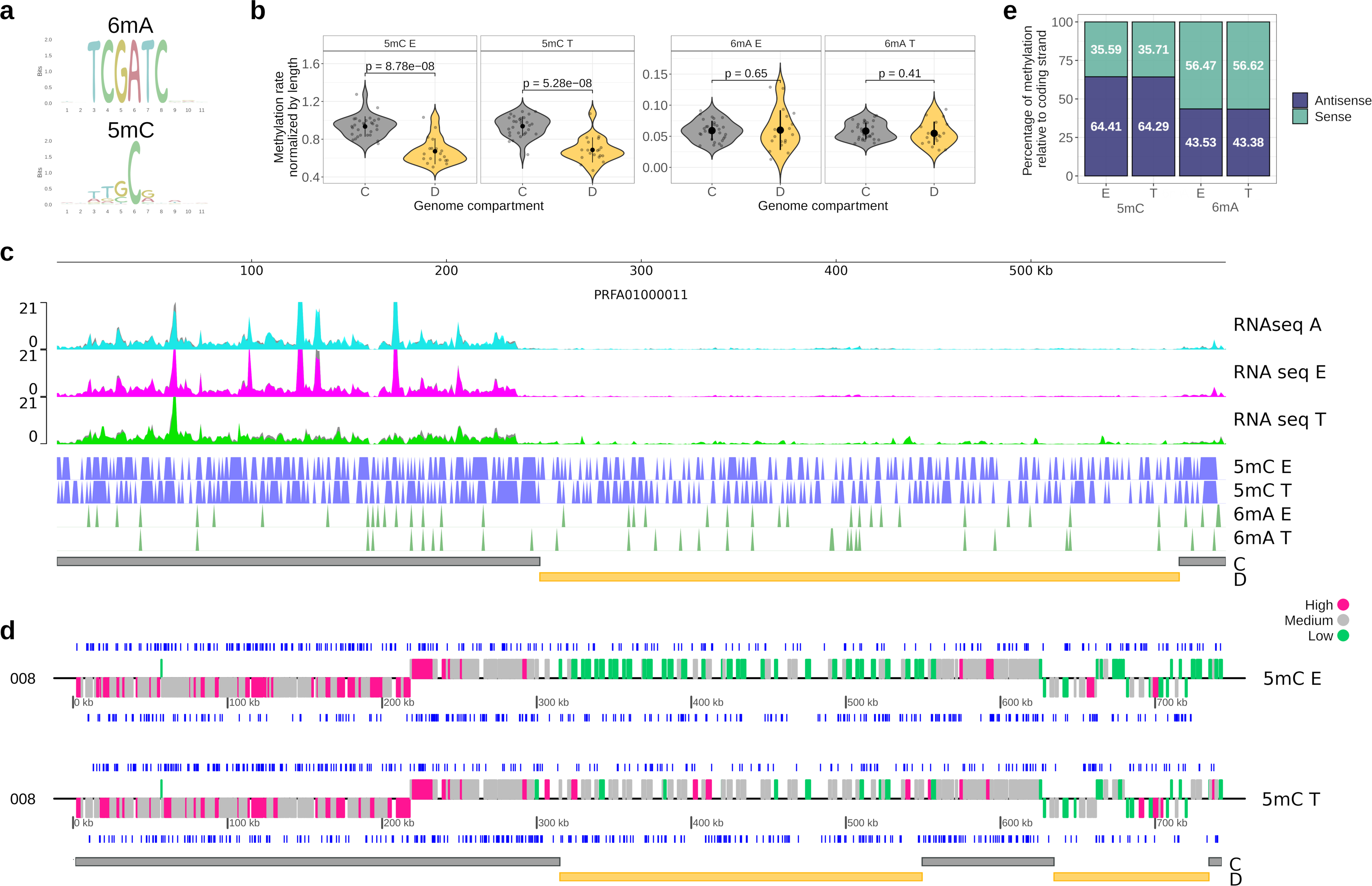
Nucleosome positioning in the genomic compartments of *T. cruzi*. **a,** Nucleosome occupancy in different regions of the genome. The difference between epimastigotes and trypomastigotes was not statistically significant (Mann-Whitney test), neither in the core nor in the disruptive compartment. **b,** Correlation between the density of well-defined nucleosomes and the percentage of disruptive genes in the scaffold. **c,** Well-defined nucleosomes in the genome compartments. Student’s *t*-test is two-sided. **d,** Representative coverage plot normalized using counts per million (CPM). Bin size 10 bp. For each condition, two biological replicates are plotted. The position of Well-defined nucleosomes in epimastigotes (E) and trypomastigotes (T) are indicated in gray. The genomic position of core (C) and disruptive (D) genome compartments are represented in grey and yellow, respectively. **e,** Nucleosome dynamics between epimastigote and trypomastigote. The NucDyn algorithm was used to detect changes (shifts, evictions, and insertions) in nucleosome architectures.

### Genome-wide identification of 5mC and 6mA in T. cruzi

DNA methylation is a crucial determinant of spatial chromatin organization and gene expression (Chen et al., 2016). Thus, we analyzed the genomic distribution of 5mC and 6mA DNA methylation marks directly detected from native DNA reads of Nanopore sequencing data in the context of the genome compartments and compared the infective and non-infective stages of *T. cruzi*. We found 6mA at low levels (0.04% of all adenines are methylated) both in epimastigote and trypomastigotes and present at the TCG6mATC motif (Supplementary Table 7 and Fig. 4a). 5mC is the predominant DNA methylation mark we found on the *T. cruzi* genome: 0.75% and 0.77% of all cytosines are methylated in epimastigotes and trypomastigotes, respectively (Supplementary Table 7). We observed a marked difference in distribution between the different compartments: the core has a higher percentage of 5mC than the disruptive on both stages (Fig. 4b and c). In addition, this modification is present mainly asymmetrically with the coding strand, with 64% of 5mC occurring in the antisense strand of coding regions (Fig. 4d and e). In concordance with little intrinsic sequence specificity around the methylated base of Eukaryotic DNA MTases, we found weakly specific motifs for 5mC. Most methylated cytosines were found in the dinucleotide GC; however, other dinucleotides were also found (Fig. 4a).

**Fig. 4:**
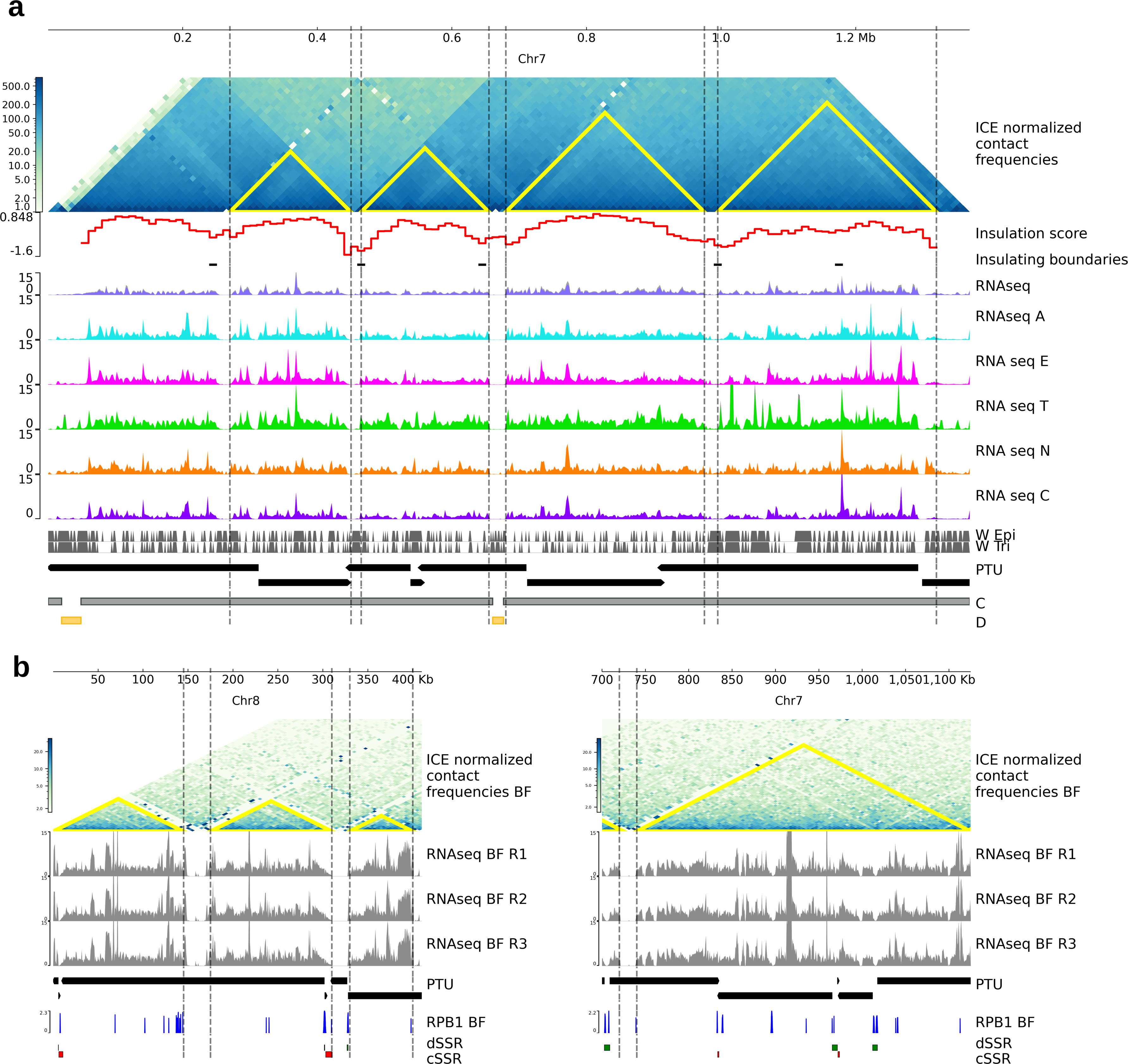
DNA methylation in *T. cruzi* genome. **a,** Sequence logo of all methylated 11-mer sequences. **b,** Analysis of DNA methylation marks in the genome compartments. Significances were determined using Unpaired two-samples Wilcoxon test (two-sided). **c,** Distribution of 5mC and 6mA marks in a representative chromosome (chromosome PRFA01000011). The genomic position of core (C) and disruptive (D) genome compartments are represented in gray and yellow, respectively. **d,** Distribution of 5mC and 6mA marks on both DNA stands in a representative chromosome (chromosome PRFA01000008). The genomic position of core (C) and disruptive (D) genome compartments are represented in gray and yellow, respectively. **e,** Quantification of DNA methylation marks relative to the coding sequence.

To obtain a better insight into the role of these epigenetic marks in gene expression regulation, we studied their genomic distribution. From the total of methylated cytosines, 50.3% are located within genes, while this number rises to 65.4% in the case of adenines. In turn, 39% (7064 of 18032) of the genes present 5mC modification, and 70% of these genes are shared between epimastigotes and trypomastigotes. Similarly, 1150 genes have the 6mA modification and half are shared between the two stages (Supplementary Table 8). Finally, we observed that many marks are conserved when parasites differentiate from epimastigotes to trypomastigotes: 38.3% and 42.7% of 5mC and 6mA marks, respectively, are shared between epimastigotes and trypomastigotes.

### The local chromatin structure determines global transcription in trypanosomes

The global expression analysis in *T. cruzi* showed that there are large stretches of the genome actively expressed that are flanked by silent regions (14kb mean length with very low or undetectable RNA levels; Supplementary Fig. S5 and Supplementary Table 9) matching in all the stages of the life cycle. Considering the gene organization and expression in trypanosomes, we expected these expressed regions to coincide with the beginning and end of DGC, constituting PTU, and those depleted of RNAs to coincide with SSR. However, we observed that most of these silent regions (65%) correspond to internal areas of the PTUs, indicating that in most cases, there is no coincidence between DGC and PTU. Strikingly, we observed that the boundaries of transcribed regions are very well-correlated with the chromatin folding domains we described (Fig. 5a). Therefore, the transcriptionally active regions of the chromosome between two silent regions are flanked by genome sites in close three-dimensional proximity. These results support that transcription is driven by the local structure of the chromatin in *T. cruzi* and that there is not necessarily a coincidence between DGC and PTU.

**Fig. 5:**
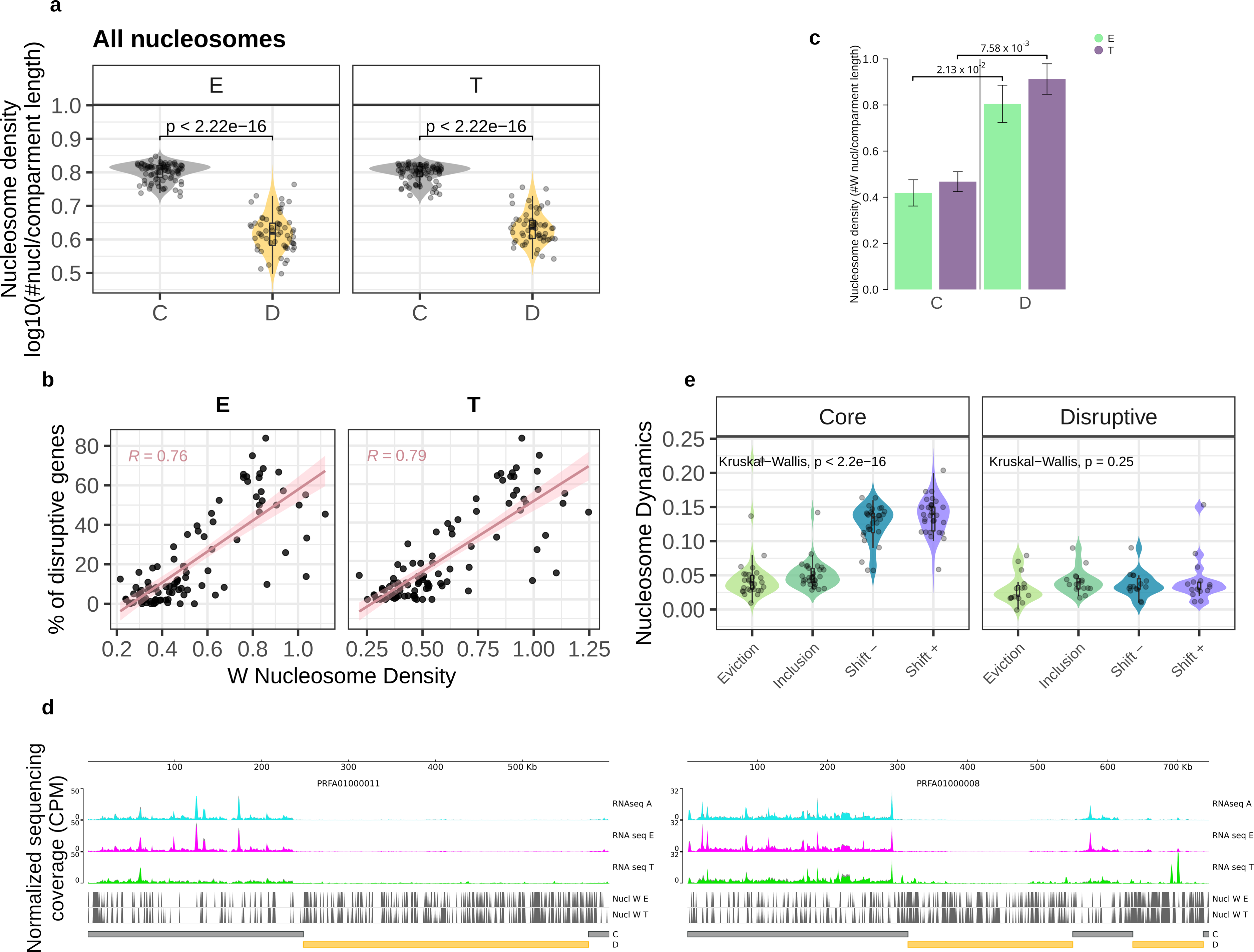
The organization of the chromatin of trypanosomes determines characteristics on gene expression. **a,** Normalized Hi-C contact map of chromosome 7 of *T. cruzi* at 10 kb resolution. CFD identified using the insulation TAD-calling algorithm from TADtool are indicated in yellow triangles. Insulation score values were calculated with FAN-C. Expression levels across the chromosome are depicted as coverage plots using a bin size of 10 bp. For each condition, two biological replicates are plotted. Vertical grey bars indicate the position of all well-defined nucleosomes. The black vertical dashed lines highlight the chromatin folding domain boundaries which align with the chromosome regions depleted for expression levels. 26% of low coverage regions correspond to SSR, 38% to CFD boundaries, 19% both to SSR and CFD boundaries and 27% to unknown reasons. **b,** Normalized Hi-C interaction frequencies of chromosomes 7 and 8 of *T. brucei* bloodstream form displayed as a two-dimensional heatmap at 5 kb resolution: chromosome 7 from 700 to 1125 Kb and chromosome 8 from 1 to 410 Kb. RNA expression across the chromosomes in the bloodstream form is depicted as normalized (counts per million, CPM) coverage plots using a bin size of 10 bp. RBP1 enrichment from the bloodstream form of *T. brucei*. Strand-switch regions (SSRs) where the transcription sense converges (cSSR) or diverges (dSSR) are indicated in red and green, respectively.

The transcriptionally active regions of the chromosome in the core compartment can span, on average, 64 kb (up to 384 Kb) and the number of genes they contain can vary from 1 to up to 173 genes (mean of 29 genes) (Supplementary Table 9). CFD boundaries suppress the expression of the genes and the transcriptionally active region in between includes fractions of different DGCs (up to three different DGCs). The presence of these regions where a large set of linearly associated genes are transcribed is not as evident in the disruptive compartment that contains the antigenic genes, even in trypomastigotes.

To study whether this is a common feature of trypanosomes, we performed the same analyses in *T. brucei*. To this end, we used Hi-C data from procyclic and bloodstream forms and RNA-seq from the bloodstream from Lister 427 strain (Faria et al., 2021; Müller et al., 2018). From Hi-C interaction maps, we performed TAD calling and observed that tridimensional genome contacts also coincide with several DGC internal regions where the transcription is interrupted in *T. brucei*. To further characterize these results, we analyzed recently published chromatin immunoprecipitation sequencing (ChIP-seq) data of the RNA pol II large subunit, RBP1 (Cordon-Obras et al., 2022). As expected, an enrichment of RBP1 was found at divergent and convergent strand switch regions where transcription starts and ends. However, the Chip-seq data also revealed a striking enrichment of RBP1 at the internal DGC regions flanked by the chromatin domains, indicating that they also constitute transcription start and end regions (Fig 5b). These results indicate that transcription initiation and termination are determined not only by DGC boundaries but also by chromatin structural domains in *T. brucei*.

### Chromosomal folding domains are conserved across strains and stages in trypanosomes

In order to determine the spatial genome organization at different stages of the *T. cruzi* life cycle and confirm that gene expression is affected by the local structure of the chromatin, we performed chromatin conformation capture (3C) experiments. To select which topological domain boundaries to analyze, we used the Hi-C maps from Brazil A4 to design 3C primers (Supplementary Table 10) in syntenic loci with Dm28c strain (Supplementary Fig. 6). The chromatin interactions we observed in Brazil A4 epimastigotes by Hi-C analysis (chr10:660000-700000 and chr29:140000-38000) were also determined in 3C assays of Dm28c epimastigotes and trypomastigotes (PRFA01000007:363000-427000 and PRFA01000013:272663-513267) (Fig. 6a and b). The resulting chromatin folding domains on Dm28c consist of 64 and 241 kb in length and are composed of 32 and 124 genes, respectively. Therefore, not only are collinearity and synteny blocks of genes at the sequence level present but also the 3D organization of the genome is maintained between the strains and different stages of the parasite life cycle.

**Fig. 6:**
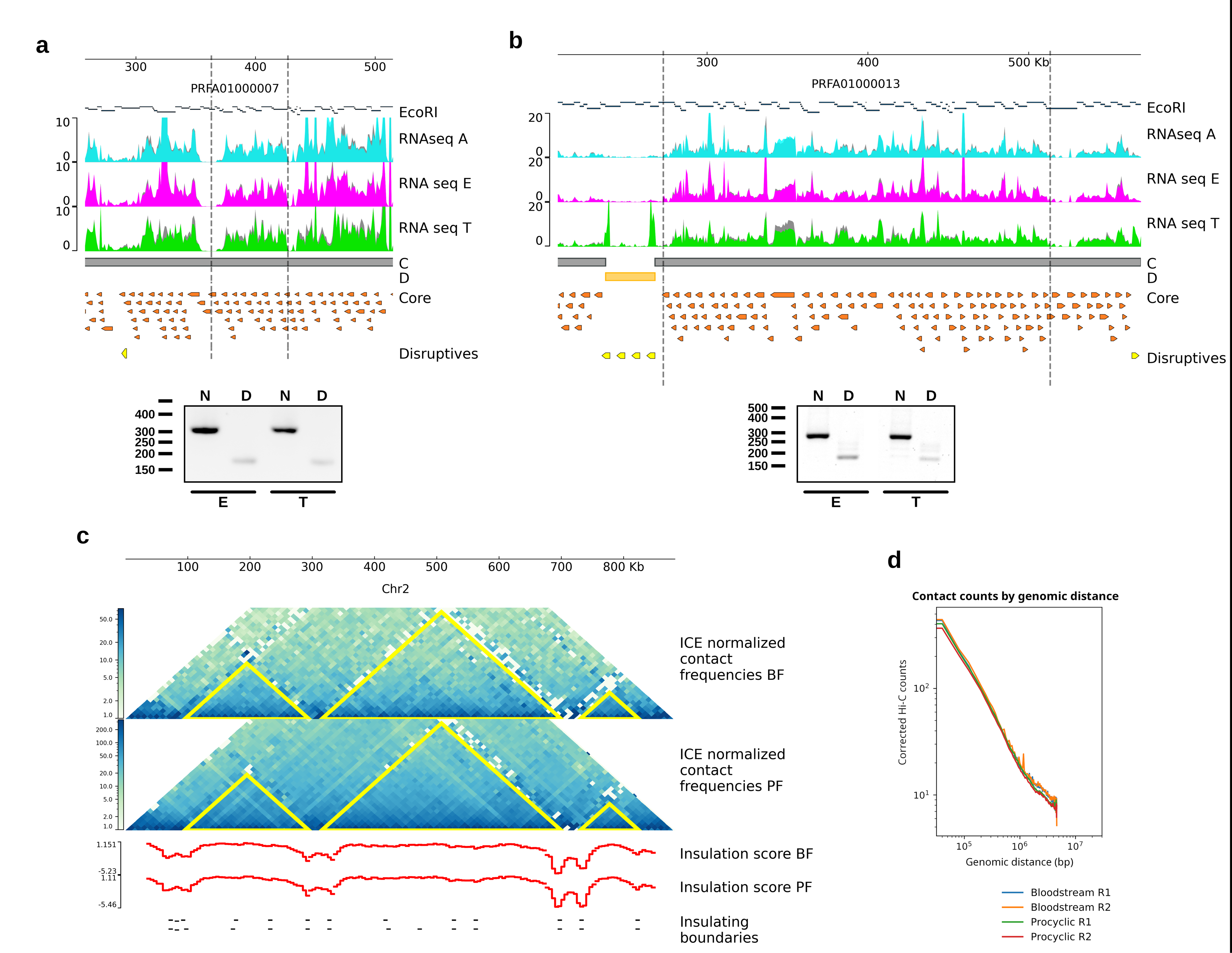
Self-interacting domain boundaries are conserved across cell types and strains. Chromosome Conformation Capture (3C) assay. PCR amplification of cross-linked DNA template cut with *Eco*RI. PCR products were separated on 2% agarose gel stained with ethidium bromide. N: neighbor Primers; D: Distance primers. **a,** Region between position 363000 and 427000 of scaffold PRFA01000007. 300 and 166 nt PCR products were amplified with N and D primers, respectively. **b,** Region between position 272663 and 513267 of scaffold PRFA01000013. 275 and 171 nt PCR products were amplified with N and D primers, respectively. **c,** Normalized Hi-C maps of procyclic and bloodstream forms at 10 kb resolution. CFD identified using TADtools are indicated in yellow triangles on Hi-C maps. The insulation score and local minima of the insulation score were calculated for both forms using FAN-C. Self-interacting chromosomal domains are conserved across African trypanosomes. **d,** We observed no differences (*p* value > 0.05) in long-range (20 kb) versus short-range contacts (<20 kb) between procyclic and bloodstream forms. BF: bloodstream form; PF: procyclic form.

Since the synteny between trypanosomes is highly conserved in the core genome, we investigated if chromatin domain conservation also occurs during different stages of *T. brucei* life cycle. In this analysis, we compared the locations of local minimums of insulation score (indicatives of regions between two self-interacting domains) identified in both stages, and we observed that most of the boundary regions are shared between procyclic and bloodstream forms (Fig. 5c). Of the 318 domain boundaries identified in the bloodstream form and 295 boundaries in procyclic form, 120 are common between both stages corresponding to 37.7% and 40.5%, respectively. Moreover, we observed no variation in the contact counts by genomic distance, neither in short-range nor in long-range contacts (Fig. 5d). These results indicate that the overall 3D organization of the core compartment of the genome is largely invariant in trypanosome genomes.

## Discussion

The genome organization of trypanosomes exhibits a common property: they are divided into a core —syntenic among trypanosomatids— and a disruptive —or non-syntenic— compartment. The latter, mainly composed of genes encoding for surface proteins, is located in the subtelomeres in the African trypanosomes (Müller et al., 2018) while is widely distributed along the genome in *T. cruzi* (Berná et al., 2018). The disruptive compartment in *T. cruzi* comprises, in several cases, almost entire chromosomes (>80%), representing a relevant difference with *T. brucei* genome organization. Although these disruptive compartments have different genomic distributions, it represents a large genome proportion in both parasites (31% of *T. cruzi* Brazil A4 genome assembled in chromosomes and 50% of *T. brucei* genome assembly (Damasceno et al., 2021), denoting the importance of these genes and the mechanisms contained in these regions for the parasites. This particular genome arrangement suggests an impact on 3D chromosome organization. In the absence of enhancer-promoter regulation, spatial chromatin conformation could constitute a relevant mechanism for regulating gene expression in these parasites. Analyzing the frequency of intra-chromosomal DNA-DNA interactions we observed that the junctions between the core and disruptive compartments are prominent insulating boundaries of chromatin interactions. Therefore, the linear genome organization (core and disruptive) has a three-dimensional —C and D— counterpart (from here, core and disruptive will refer to the linear organization of the trypanosome genome, and C and D to their three-dimensional organization). Equivalent to the C and D in *T. cruzi*, the core and subtelomeric regions were previously defined as 3D compartments in *T. brucei* (Müller et al., 2018).

We investigated the juxtaposition of distal chromosomal loci in the nuclear space of *T. cruzi,* and we found that the genome is organized into self-interacting chromosomal domains. We referred to them as CFD since they are significantly smaller than eukaryotic TADs, as expected in organisms where most chromosomes are shorter than 3 Mb, and exhibit reduced non-coding regions (e.g., lack of introns, short intergenic and inter DGC regions). The CFDs are smaller in the D compartment (almost half the size as those in the core) and present a higher frequency of contacts, which indicates that chromatin is more compact in this compartment, which is in agreement with recent FAIRE-seq analyses (Lima et al., 2022). Notably, while the core compartment interactions are mainly intra-chromosomal, the disruptive compartment exhibits a high frequency of inter-chromosomal contacts. This higher compaction of D chromatin in trypanosomes may have both the role of avoiding spurious transcription and also facilitating DNA recombination events. TS, TcMUC, and MASP proteins cover the surface coat of *T. cruzi*; hence, they are exposed to the immune system. Members of the TS multigene family are subject to frequent recombination (Weatherly et al., 2016), and the number of pseudogenes probably represents traces of past recombination events and/or reservoirs for future events. TcMUC and MASP multigene families exhibit the same characteristics, although their specific functions have yet to be well known. The close three-dimensional proximity of these multigene families combined with their high sequence similarity constitutes a scenario favorable for stochastic recombination events, generating antigenic variability and/or relocation of genes favoring their up- or down-regulation, essential for the evasion strategy of the immune system of *T. cruzi*. Comparable, in *T. brucei* the global mechanism implies the recombination of the VSG genes into subtelomeric regions to switch the expression of the protein that covers the parasite’s surface as the infection progresses and the immune system matures its response (McCulloch et al., 2015). Therefore, both parasite genomes exhibit a D compartment composed of species- and stage-specific surface proteins, which would generate favorable conditions for generating antigenic diversity, either by recombination, allelic exclusion, or differential transcription. On the contrary, it is expected that the core compartment, which does not undergo antigenic variation, presents less variability concerning the structure of the chromatin. In agreement with this, we observed well-conserved CFDs in this C compartment in the different stages and strains in both parasites. Thus, the main differences in genome architecture are those that allow *T. brucei* to survive extracellularly and *T. cruzi* to adapt to both intracellular and extracellular forms.

One mechanism to ensure coordinated regulation of gene expression of large stretches is the formation of higher-order 3D chromatin structures (Bhat et al., 2021). Given that we observed that chromatin is differently organized in C and D compartments, we investigated whether this three-dimensional genome organization implies selective regulation of gene expression. We found that the core compartment is mainly composed of genes with high and medium expression, while most of the genes from the disruptive compartment exhibit lower expression. The core functions as a constitutively expressed region since its genes are expressed similarly throughout the life cycle. Conversely, 80% of the DEG upregulated in trypomastigotes are disruptive genes, which implies that the disruptive compartment constitutes a specialized region for the infective stage. Moreover, disruptive genes present lower overall expression values, and a discrete number of genes highly increase their expression in trypomastigotes. These findings modify the accepted paradigm that these parasites transcribe indiscriminately and then modulate post-transcriptionally their expression. This is clearly observed in the core compartment, but it is not applicable to the disruptive compartment since very few of the hundreds of genes in each multigenic family are significantly expressed. Both in *T. cruzi* and *T. brucei*, the disruptive compartment constitutes regions of high chromatin compaction, which could favor gene-to-gene regulation, preventing spurious transcription and favoring intra- and inter-chromosomal recombination. In *T. brucei*, multiple VSG genes are transcribed before a single gene is chosen (Hutchinson et al., 2021) and this choice relies on the formation of an extra-nucleolar structure named expression-site body that contains a local reservoir of RNA polymerase I and RNA processing factors (Navarro et al., 2001; Faria et al., 2021). In *T. cruzi* the expression of the highly expressed surface genes is probably determined by the nucleoplasmic location of the loci and the proximity to the transcription and processing machinery, but HiC data of the infectious stage is needed to study this and other aspects of genome biology at this stage of the life cycle.

The analysis of RNA-seq data revealed a close interrelationship between chromatin domains and RNA levels in the core compartment. The genome regions between two CFDs are characteristically RNA silent, while the regions contained in CFDs are actively transcribed, indicating that the expression is strongly determined by the 3D organization of the chromatin and does not necessarily correlate with the DGC. Therefore, the paradigm that there is a correspondence between DGC and PTU is not always fulfilled. Importantly, this is more than just a feature of *T. cruzi*. In *T. brucei*, when we eliminated the bias of DGC as a reference for transcription start and end, we also found a similar correlation between CFD and transcription in the core compartment, denoting a common feature of trypanosomes. RBP1 is enriched in these DGC-internal regions that are flanked by CFDs, endorsing these are regions where transcription starts and ends. Future studies are necessary to determine whether these DGC-internal transcription start and stop sites present similar characteristics as those previously described in the SSR (e.g., histone modifications, histone variants, J base, methylation). Taken together, these results strongly support that the flow of genetic information from DNA to RNA in trypanosomes is regulated at transcription initiation through epigenetic mechanisms.

In order to determine other epigenetic mechanisms that may affect gene expression, we analyzed the organization of the nucleosomes and the global DNA methylation. Nucleosome positioning studies have revealed that the disruptive compartment is generally densely packed with well-defined nucleosomes, suggesting a less permissive chromatin structure. This is consistent with a more condensed nature and in agreement with previous reports showing that in the insect stages of *T. cruzi* the disruptive compartment has less open chromatin than the core (Lima et al., 2022). DNA methylation marks are also differentially distributed in the genome compartments. We found that 5mC is more predominant than 6mA in *T. cruzi* genome and presents an asymmetric distribution between compartments: there is more 5mC mark in the core than in the disruptive compartment. 6mA was nearly undetectable, and we found no difference in the abundance of 6mA levels among the compartments. However, the sequence specificity denotes the presence of a specific DNA methylase for introducing 6mA. It is increasingly clear that the relationship between DNA methylation and transcription is complex and highly dependent on the context (de Mendoza et al., 2022). In turn, in *T. cruzi*, where there is not one promoter per gene, the situation may differ from what generally occurs in other eukaryotes. The fact that 40% of the marks are preserved between trypomastigotes and epimastigotes means that they are maintained after DNA replication and differentiation, positioning them as proper epigenetic marks.

The most accepted idea regarding the differential regulation of gene expression in trypanosomes is that transcription is constitutive and post-transcriptional mechanisms individually modulate that transcript levels. However, here we provide evidence that the expression of the disruptive compartment in *T. cruzi* does not respond to this widely accepted model: most of their genes exhibit low and undetectable levels, and a discrete number greatly increases their expression in the trypomastigote stage. These results do not go against the statement “trypanosomes mainly regulate at the post-transcription level”, but rather this would be the expression modulation model in the core compartment. We propose that there are two major regulation models and that they are spatially isolated in the nucleoplasm by the three-dimensional C and D compartments in which the chromatin is organized. The compartmentalization of the genome makes it possible to isolate the molecular mechanisms necessary for regulating each type of gene. We propose that the compartments previously defined by the linear association of genes are functional compartments for the regulation of gene expression and genome plasticity. Incorporating time-course data and more data types will reveal further insights into the mechanisms of 3D chromatin structure and gene regulation in trypanosomes. Single-cell studies will better explain how antigenic variability occurs in *T. cruzi*, describing how highly expressed disruptive genes change with the course of infection and how the expression is in individual parasites.

## Methods

### Cell culture

*T. cruzi* Dm28c (Contreras et al., 1988) parasites were cultured axenically in liver infusion tryptose medium supplemented with 10% (v/v) inactivated fetal bovine serum (*GIBCO*) at 28°C. Trypomastigotes were collected from infected Vero cell line monolayers’ supernatant and maintained cyclically. Vero cells were cultivated in Dulbecco’s Modified Eagle’s Medium [DMEM(1×) + GlutaMAX™-l, Gibco® by Life Technologies™] supplemented with 10% (v/v) fetal bovine serum (FBS, *GIBCO*), penicillin 100 U/mL and 100 µg/mL streptomycin (*Thermo SCIENTIFIC*) at 37°C in a humidified 5% CO2 atmosphere.

### Methylation analysis

As previously described (Díaz-Viraqué et al., 2021), high molecular weight genomic DNA isolation was performed using phenol/chloroform/isoamyl alcohol extraction followed by EtOH precipitation. Purified gDNA was fragmented to 10 kb using g-TUBEs (*Covaris*) to maximize the yield of nanopore sequencing. Genomic libraries were prepared using the Ligation Sequencing Kit (SQK-LSK109) and Native Barcoding Expansion Kit (EXP-NBD104) (Oxford Nanopore Technologies, United Kingdom) following the protocol Native barcoding genomic DNA. Pooled samples were quantified using Qubit dsDNA HS Assay Kit, loaded onto an R9.4.1 Flow Cell (FLO-MIN106D) and sequenced on an Oxford Nanopore Technologies (ONT) MinION platform for 72 h. The coverage per sample ranged from 29X to 34X. Raw data (multi_read_fast5 files) were converted to single_read_fast5 files using ont_fast5_api v3.1.6 (ONT) and basecalled by using Guppy v3.6.0 (ONT) with dna_r9.4.1_450bps_modbases_dam-dcm-cpg_hac.cfg configuration. The strand-sensitive and single-nucleotide-based detection of DNA base modifications was performed with DeepMod (Lui et al., 2019), which detects m5C and m6A in DNA using recurrent neural networks (RNN) model. Only genomic positions with >5 read coverage and >90% of methylation percentage were considered for the analysis. Dm28c reference genome was used as input and methylation positions were assigned to different genomic regions using bedtools v2.27.1 (Quinlan et al., 2010). Core and disruptive methylation quantifications were made in the largest 30 scaffolds of the Dm28c 2018 genome assembly, representing 38% of the genome size.

### Nucleosome position analysis

MNase-seq data was obtained from NCBI (BioProject PRJNA665060) (Lima et al., 2021). Quality control was performed using FastQC, and reads with less than 70 bp after trim low-quality nucleotides (--nextseq-trim=20) and clip the adapter sequence were removed using cutadapt (Martin et al., 2011). For nucleosome calling, paired-end reads were aligned to both reference genomes Dm28c (TcruziDm28c2018 from TriTrypDB version 51) and Brazil A4 (TcruziBrazilA4 from TriTrypDB version 53) using Bowtie v1.3.1 (Langmead et al., 2009) allowing up to 2 mismatches and maximum insert size of 500 bp. The resulting BAM files were imported and processed in R in order to merge biological replicates by chromosome. After duplicate removal with Picards MarkDuplicates (Picard Toolkit. 2019), the Fourier transform filtering to remove noise and peak calling to accurately define and classify the location of nucleosomes across the genome were performed with nucleR (Flores et al., 2011). Well-positioned nucleosomes were considered when nucleR peak width score and height score were higher than 0.6 and 0.4, respectively. Finally, nucleosome dynamics (Buitrago et al., 2019) was used to study differences between stages: occupancy differences (insertions and evictions) and displacement of nucleosomes (shifts).

### Chromatin Conformation Capture 3C

Parasites (1 x 10^8^ trypomastigotes and 5 x 10^9^ epimastigotes) were collected by centrifugation at 2200×g for 10 min at room temperature and washed twice with PBS 1X. Cells were fixed with 1% formaldehyde for 15 min at room temperature on a shaker. The cross-linking was quenched by adding 2.5 mL of 1 M glycine (125 mM) and incubated for 5 min at room temperature, then on ice for 15 min. Fixed cells were then centrifuged, washed twice with ice-cold PBS 1X, and resuspended in 1.4 mL of cold lysis buffer (10 mM Tris·HCl pH 8, 10 mM NaCl, 0.2% Nonidet P-40, 1× Protease Inhibitors). After homogenization with a Dounce homogenizer pestle A, the nuclei were washed twice with 1.25× rCutSmart™ Buffer (New England BioLabs) and resuspended in 0.5 mL of rCutSmart™ Buffer (New England BioLabs) containing 0.3% SDS. Samples were incubated for 40 min at 65 °C and then for 20 min at 37 °C. 2% Triton X-100 was added and the nuclei were further incubated for 1 h at 37°C to sequester the SDS. Cross-linked DNA was digested using 400 U of EcoRI-HF® (R3101 New England BioLabs® Inc) overnight at 37 °C and the enzyme was inactivated with 1% SDS at 65 °C for 20 min. Digestion products were diluted by the addition of 4 mL 1.1× T4 DNA Ligase Reaction Buffer (New England BioLabs® Inc) with 1% Triton X-100 and incubated for 1 h at 37°C. DNA was ligated for 4 h in a water bath at 16 °C (30 Weiss Units of T4 DNA Ligase, M0202 New England BioLabs® Inc). 300 μg of Proteinase K was used to reverse the cross-linking overnight at 65°C. Finally, samples were incubated with 300 μg RNase A, and the DNA was purified by phenol extraction followed by ethanol precipitation. The Control library with all possible ligation products was prepared in parallel using non-crosslinked genomic DNA. Ligation products were detected and quantified in control and cross-linked libraries by PCR using *locus*-specific primers. We designed a set of 6 oligos excluding sequences from repetitive regions (Supplementary Table 10). The linear range of amplification was determined for all the libraries by serial dilution of DNA amounts.

### Transcriptomic analysis

Total RNA from cell-derived trypomastigotes, epimastigotes, and intracellular amastigotes were treated with RiboZero Gold magnetic beads - eukaryotic kit (250ng as input) as described previously (Greif et al., 2019). Library preparation of purified RNA was then performed using TruSeq Stranded Total RNA kit (Illumina) with random primers. 80 bp paired-end reads were generated using an Illumina NextSeq 500 MID platform. Raw reads were processed with cutadapt (Martin et al., 2011) to trim low-quality nucleotides (--nextseq-trim=20) and clip the adapter sequence. The resulting reads with less than 70 bp were discarded (--minimum-length=70). Trimmed reads were checked for per-base quality using FastQC. Cleaned reads were then aligned to the Dm28c reference genome (TcruziDm28c2018 from TriTrypDB version 51) using Hisat2 (--no-spliced-alignment --rna-strandness RF) (Kim et al., 2015) and mapped reads with a mapping quality score (MapQ) <10 were discarded with SAMtools as it was previously described (Hutchinson et al., 2016). The coverage files were smoothed, and counts per million (CPM) normalized in a bin size of 10 bp using bamCoverage function from deeptools (Ramírez et al., 2016). Salmon (Patro et al., 2017) was used to estimate transcript levels and statistical analysis of differential expression of mRNAs was tested with DESeq2 (Love et al., 2014). The expression of each transcript was quantified in transcript per million (TPM) units. Consistency between replicates was assessed with Pearson correlation and principal component analysis (PCA). Genes were considered as differentially expressed when they were statistically significant (padj < 0.001) and had a fold change in transcript abundance of at least two (|log2FC| > 1). To study the dynamic of RNA levels in genome compartments along the stages, we classified the transcripts into three groups based on their relative abundance. High, medium, and low expression genes were determined using quantiles on normalized counts. Also, to investigate expression patterns, we classified the scaffolds into “core”, “disruptive”, and “mixed” according to the percentage of compartments that make them up. The transcriptomic data corresponding to *T. cruzi* different subcellular compartments, Brazil A4 and *T. brucei* were analyzed identically.

To study immature/nascent transcripts, 5′ UTR regions were first annotated and assigned to protein-coding genes using UTRme (Radío et al, 2018). SLA sites identified at the same positions in all transcriptomes were used for the analysis to avoid bias of differential processing between stages. Alignments of reads spanning SLA sites were quantified, normalized, and classified as processed and unprocessed according to the presence or absence of spliced leader sequence in the read, respectively.

To identify RNA zero coverage regions, genome coverage for all positions in BEDGRAPH format was calculated from mapping quality filtered bam files using genomecov function from bedtools v2.27.1 (Quinlan et al., 2010) with -bga option. Then, all regions of the genome with zero coverage were extracted, and intervals with <500 bp length were filtered out.

### Chromatin immunoprecipitation analysis

The 37 x 2 bp paired-end raw reads were processed with cutadapt (Martin et al., 2011) to remove low-quality bases from both ends (-q 30), remove flanking N bases from each read (--trim-n) and discard reads containing more than 5 N bases (--max-n 5). The resulting reads with less than 35 bp were discarded (--minimum-length=35). Raw and trimmed reads were checked for per-base quality using FastQC. Cleaned reads were then aligned to *T. brucei* reference genome (TbruceiLister427_2018 from TriTrypDB version 53) using Bowtie v1.3.1 (Langmead et al., 2009) allowing two mismatches (-v 2). Unique alignments reported (-m 1) were compressed and sorted using SAMtools (Li et al., 2009), and duplicate reads were removed using Picard (MarkDuplicates). Normalization and peaking calling was conducted using MACS2 (Zhang et al., 2008) with a false discovery rate of 5% and filtering out peaks with low fold-enrichment (-q 0.05 and --fe-cutoff 1.5). The –g parameter was set at 50081021.

### Chromatin Conformation Computational analysis

Hi-C data sets were processed from raw reads to normalized contact maps using HiC-Pro v3.1.0 (Servant et al., 2015). Quality as assessed using MultiQC. TAD calling was performed with TADtools (Kruse et al., 2016), hicFindTADs function of HiCExplorer v3.7.2 (Ramírez et al., 2018). Insulation score was calculated at 30, 50, 70, 100 and 150 kb sliding windows with FAN-C (Kruse et al., 2020). The data integration, comparison, and visualization of the different data sets were performed using pyGenomeTracks (Lopez-Delisle et al., 2020).

**Supplementary Fig. S1: Normalized Hi-C interaction frequencies of chromosome 3, 9 and 10.** Results are displayed as a two-dimensional heatmap at 10 kb resolution. Chromatin Folding Domains (CFDs) identified using HiCExplorer and TADtools (triangles on Hi-C map) are indicated in yellow. The local minimum of insulation score calculated using FAN-C (representing the region between two self-interacting domains) are indicated as black dash. The genomic position of core and disruptive genome compartments are represented in gray and yellow, respectively. Hi-C coverage is plotted.

**Supplementary Fig. S2: Representative RNA-seq sequencing coverage plots in entire scaffolds normalized using counts per million (CPM).** Bin size 10 bp. For each condition (A: amastigotes; E: trypomastigotes; T: trypomastigotes), two biological replicates are plotted.

Supplementary Fig. S3: Genomic distribution of genes classified according to RNA expression (RNA-seq) in *T. cruzi* Brazil A4 strain. Chromosomes of Brazil A4 genome assembly. Disruptive genes (TS, TcMUC and MASP) are indicated in black.

**Supplementary Fig. S4: Genomic distribution of genes classified according to the expression level (RNA-seq) in nucleus and cytoplasm.** Disruptive genes (TS, TcMUC and MASP) are indicated in black.

**Supplementary Fig. S5: Data integration.** Normalized Hi-C interaction frequencies of chromosome 10 of *T. cruzi* displayed as a two-dimensional heatmap at 10 kb resolution. Chromatin Folding Domains (CFDs) identified using TADtools are indicated as triangles on the Hi-C map. Insulation score values were calculated using FAN-C and the black horizontal dashes represent the local minima of the insulation score, indicatives of the domain boundaries. RNA expression across the chromosome is depicted as coverage plots using a bin size of 10 bp. For each condition, two biological replicates are plotted. A: amastigotes; E: epimastigote; and T: tripomastigote forms. Vertical gray bars indicate the position of all well-defined nucleosomes. Polycistronic Transcription Units (PTUs). This is a core chromosome as the core compartment comprises the entire chromosome. Some disruptive genes present in this chromosome are indicated as yellow arrows. Zero coverage regions (≥500 bp length) are shown as black vertical lines. Low coverage regions were determined considering all transcriptomes and are represented as blue blocks.

**Supplementary Fig. 6: Whole chromosomal alignments.** Chromosome 10 from *T. cruzi* Brazil A4 presents synteny with two scaffolds of *T. cruzi* Dm28c genome assembly. The region we studied is present in scaffold PRFA01000007. Chromosome 29 from *T. cruzi* Brazil A4 presents synteny with scaffold PRFA01000013 of *T. cruzi* Dm28c genome assembly.

**Supplementary Table 1: Classification of the *T. cruzi* chromosomes.** *T. cruzi* Dm28c and Brazil A4 chromosomes were classified as Core, Disruptive or Mixed according to the genome compartment composition. Chromosomes were classified as core or disruptive if one of these genomic compartments covers more than 80% of the length of the chromosome, while the rest are considered Mixed.

**Supplementary Table 2: Chromatin folding domains.** CFD predictions of *T. cruzi* using different algorithms.

**Supplementary Table 3: Genome features at chromatin domain boundaries.** Identification of genomic features at the CFD boundaries.

Supplementary Table 4: Differentially expressed genes along the life cycle stages of *T. cruzi*. Result of differential gene expression analysis based on the negative binomial distribution.

**Supplementary Table 5: Normalized read counts.** High, medium and low expression genes were determined using quantiles on normalized counts (transcript per million, TPM). It is also indicated if the gene is DEG in the comparison of epimastigotes vs trypomastigotes.

Supplementary Table 6: Nucleosome positions in *T. cruzi* genome. Nucleosomes identified and classified using nucleR.

**Supplementary Table 7:** Deepmod

Supplementary Table 8: DNA methylation in *T. cruzi*.

**Supplementary Table 9: Length of low or zero coverage regions on core chromosomes.** As the zero cov regions in all transcriptomes are difficult to define in the disruptive compartment because it is a stage-specific expression compartment, The analysis of these regions was carried out on chromosomes defined as core chromosomes (80-100% of chromosome length correspond to core compartment): Chr1, Chr4, Chr7, Chr8, Chr10, Chr14, Chr15, Chr16, Chr17, Chr18, Chr22, Chr23, Chr28.

**Supplementary Table 10: Chromatin Conformation Capture 3C primers.**

## Data availability

Raw sequencing data for Illumina and Oxford Nanopore Technologies platforms were deposited to NCBI: RNA-seq to BioProject ID PRJNA850400 and Mehtylation to BioProject ID PRJNA935260. Hi-C and RNA-seq data for T. cruzi Brazil_A4 epimastigotes from a previous study (Wang et al., 2021) are available from the NCBI under accession number SRR11803985 and SRR12792489. Hi-C data for T. brucei PF and BF, as well as RNA-seq data from BF from previous studies (Faria et al., 2021; Müller et al., 2018) are available from the NCBI SRA under accession numbers: ERR3712002, ERR3712009, SRR7721317, SRR7721318 and SRR5809498-SRR5809500. RBP1 Chip-Seq (Cordon-Obras et al., 2022) and MNaseq (Lima et al., 2021) data are available under SRR9022833, SRR9022834, SRR13260277, SRR13260279 and SRR12710803-SRR12710808 accession numbers. RNA-seq data from different subcellular compartments of T. cruzi (Pastro et al., 2017) are available from the NCBI under accession number SRR4232036-SRR4232038.

## Acknowledgments

We thank Anton Enright (University of Cambridge, UK) for facilitating the RNA-seq sequencing; Nathaniel G. Jones (University of York, UK) for critical reading of the manuscript; and Ayelen Lizarraga (University of Chicago) and other members of the Laboratorio de Interacciones Hospedero-Patógeno-UBM (Institut Pasteur de Montevideo) for their helpful comments and many interesting discussions.

