## Supplementary figures and images for "3D genome organization drives gene expression in trypanosomes"

### Supplementary_Fig1.pdf

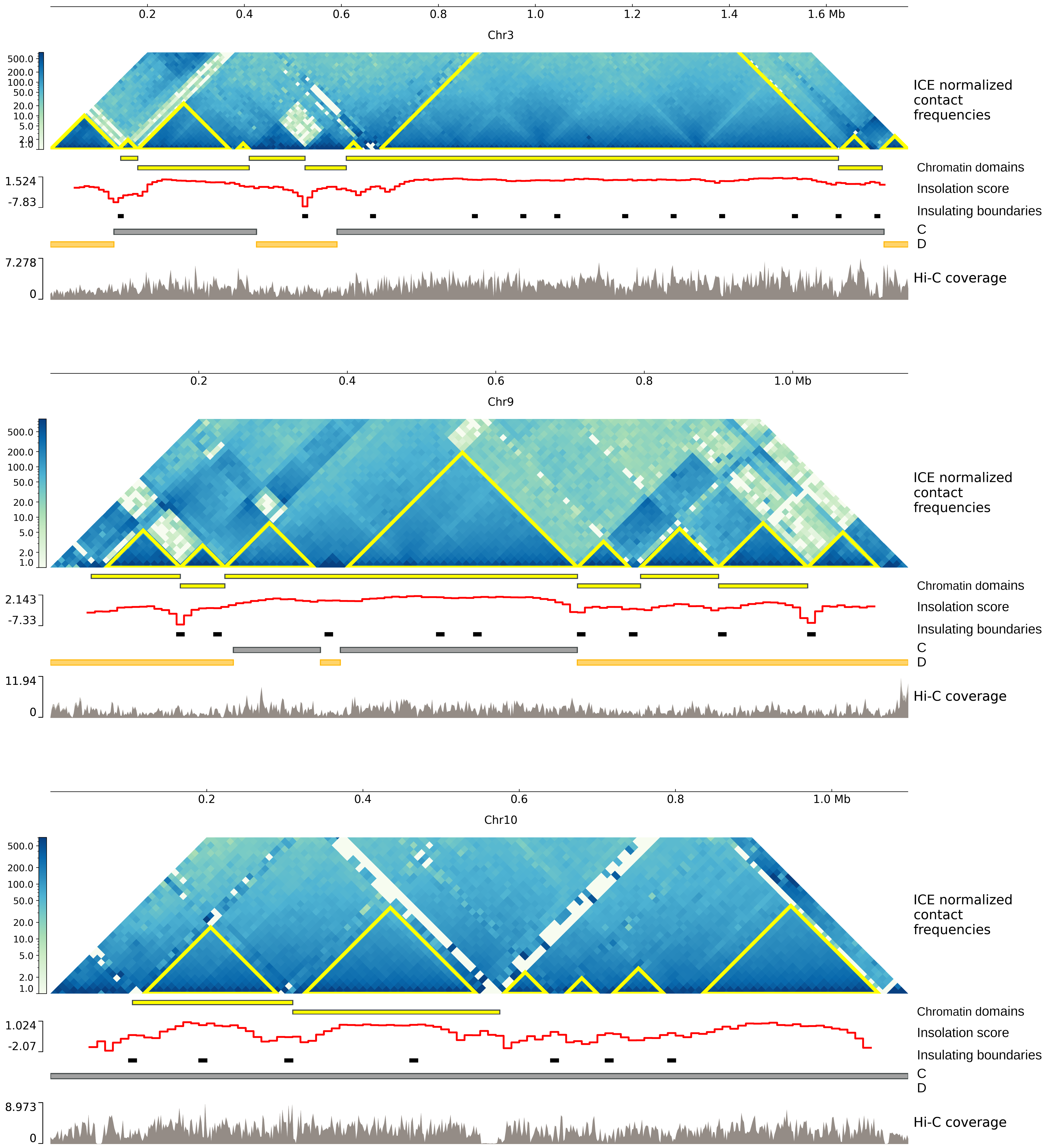
